# Mirages in continuous directed enzyme evolution: A cautionary case study with plantized bacterial THI4 enzymes

**DOI:** 10.1101/2024.11.19.624164

**Authors:** Kristen Van Gelder, Anuran K. Gayen, Andrew D. Hanson

## Abstract

Continuous directed evolution (CDE) improves enzyme characteristics by hypermutating the enzyme gene *in vivo* in a microbial platform, linking enzyme activity to growth, and selecting for growth rate. Combined with genome editing, CDE can expand the gene pool for plant breeding. THI4 enzymes, essential for thiamin synthesis, are ideal targets for plant CDE. Plant THI4s are inefficient; their replacement by efficient bacterial THI4s could boost biomass yield by up to 4%. However, bacterial THI4s are O_2_-sensitive and unsuited to plants. Previous CDE campaigns in the yeast OrthoRep system adapted bacterial THI4s for plant-like conditions, achieving success with *Mucinivorans hirudinis* THI4 (MhTHI4), which acquired beneficial mutations that improved growth. Here, we increased selection pressure on MhTHI4 by reducing its expression, leading to faster-growing populations with new nonsynonymous and synonymous mutations. Surprisingly, the synonymous mutations appeared largely responsible for the growth rate improvements, providing a cautionary example for other OrthoRep CDE projects.

---

Continuous directed evolution (CDE) improves the characteristics of a target enzyme by hypermutating the enzyme gene *in vivo*, coupling enzyme activity to growth of a microbial platform, and selecting for growth rate (Molina *et al*., 2022). Directed evolution can be interfaced with genome editing to expand the gene pool available for plant breeding; this powerful combination (DE-GE) has been neatly termed ‘a Green (r)Evolution’ (Gionfriddo *et al*., 2019). THI4 enzymes, which make the thiazole moiety of thiamin, are good testbed targets for plant CDE technology. Plant THI4s are energy-inefficient suicide enzymes that could potentially be replaced by efficient, non-suicide bacterial THI4s to increase biomass yield by as much as 4% (Joshi *et al*., 2021). However, bacterial THI4s are O_2_-sensitive and otherwise ill-adapted to plants (Joshi *et al*., 2021). We therefore previously ran CDE campaigns in the yeast OrthoRep system to ‘plantize’ bacterial THI4s, i.e., to improve function in an aerobic, plant-like milieu (Figure **1a**) (García-García *et al*., 2022). Two notably successful campaigns were for the THI4 from *Mucinivorans hirudinis* (MhTHI4); these campaigns culminated when populations acquired single V124A or Y122C mutations that improved growth to near the wildtype rate (Van Gelder *et al*., 2023). Such culmination can be overcome by increasing the selection pressure (Molina *et al*., 2022).

**Figure 1.**
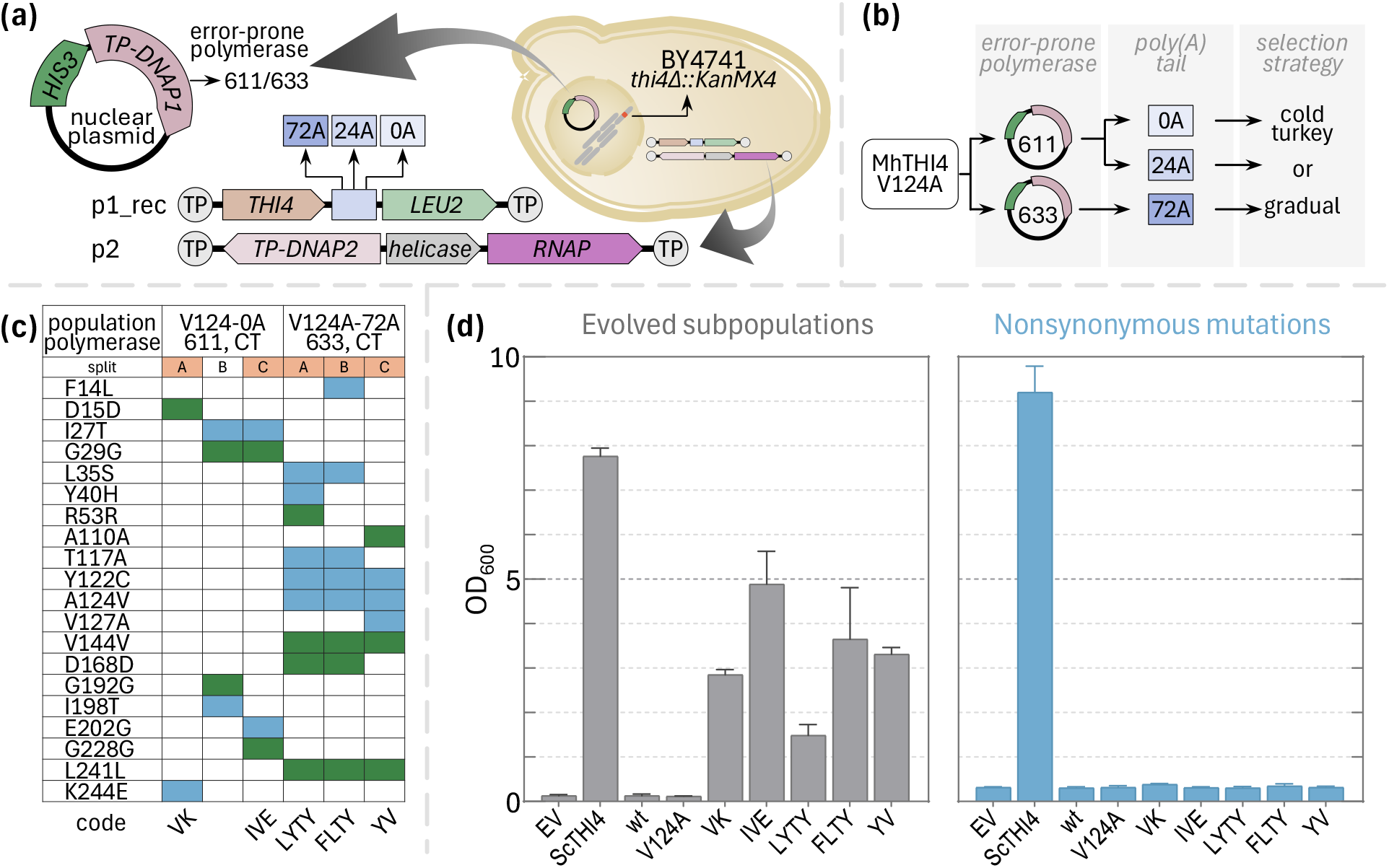
OrthoRep campaigns to plantize a bacterial THI4 and their outcomes. (a) The OrthoRep system. The target enzyme (MhTHI4 V124A), plus or minus a poly(A) tail, is encoded on the cytoplasmic p1 plasmid that also carries a LEU2 marker. p1 is hypermutated by a p1-specific, error-prone DNA polymerase (TP-DNAP1_611 or TP-DNAP1_633) encoded on a nuclear plasmid. The BY4741 platform strain carries a *thi4*Δ deletion to couple growth to the activity of the THI4 on p1. (b) The combinations of expression-reduction regimes with cold turkey (CT) or gradual (G) selection. (c) The nonsynonymous (blue) and synonymous (green) mutations that had swept populations by the end of campaigns. When populations failed to grow early in campaigns, surviving populations were split into subpopulations (A, B, C) and propagated independently. The five mutant sequences whose nonsynonymous mutations were tested are indicated in orange. (d) Growth of the five evolved populations from which these mutant sequences came compared to the lack of growth supported by these sequences when purged of synonymous mutations and cloned into fresh p1 and fresh cells (with a 72A tail and TP-DNAP1_611). The parent V124A mutant and yeast THI4 (ScTHI4) were included as benchmarks as well as an empty vector (EV) control. Data are mean ± SE for the last three passages of the campaigns for evolved subpopulations and for 12 replicate cultures for the tests of nonsynonymous mutations.

In the present work, we increased selection pressure on the MhTHI4 target by reducing its expression, thus reducing enzyme activity and cell growth rate and renewing scope for improvement. In OrthoRep, expression can be reduced by shortening the genetically-encoded poly(A) tail of the target’s mRNA or reducing the copy number of the plasmid (p1) bearing the target gene (Ravikumar *et al*., 2018; Zhong *et al*., 2018) (Figure **1a**). These maneuvers led to improved, i.e., faster-growing, populations harboring MhTHI4s with new nonsynonymous mutations, plus synonymous ones. Surprisingly, testing indicated that the synonymous mutations were probably largely or wholly responsible for the observed improvements in growth rate. As such ‘mirages’ seem highly likely to appear in other OrthoRep CDE projects, we document them here as a cautionary case study.

Our previous campaigns with MhTHI4 used a commercially recoded gene, of which the V124A mutant (Van Gelder *et al*., 2023) was the starting point for the present campaigns (Appendix **S1**). As before, the platform strain was BY4741 *thi4*Δ, which requires a thiazole precursor (HET) or thiamin for growth. The poly(A) tail was shortened from ∼72A to 24A or 0A to reduce expression twofold or tenfold, respectively (Zhong *et al*., 2018). To reduce p1 copy number, the error-prone DNA polymerase was changed from TP-DNAP1_611 to TP-DNAP1_633, which also raises the mutation rate (García-García *et al*., 2022). Each expression-reduction regime was combined with ‘cold turkey’ selection (culture without added thiamin or HET) or ‘gradual’ selection (initial supplementation with limiting thiamin or HET that is tapered to zero) (García-García *et al*., 2022). Subculturing was every 4-6 days. Nine independent populations of the V124A mutant were engineered for each expression-reduction regime and subjected to cold turkey or gradual selection, giving 54 initial subpopulations (Figure **1b**). Subpopulations that survived without supplementation were split in three (denoted A, B, C) and propagated independently. Sequencing bulk DNA from subpopulations identified mutations of interest, i.e., those that had fully replaced a wildtype base (‘swept’ the subpopulation).

The cold turkey strategy proved effective; after 33 passages it yielded two subpopulations (V124A with no A tail and V124A with TP-DNAP_633) whose splits all reached OD_600_ 1.5–5.0 by the end of each passage. All told, eleven nonsynonymous mutations were found in the selected subpopulations along with nine synonymous mutations (and two 10B2 promoter mutations, most likely neutral because 10B2 is already optimized; Zhong *et al*., 2018) (Figure **1c**). Notable nonsynonymous mutations were a reversion of V124A and its displacement by Y122C; this switch implies that Y122C is functionally superior.

Five sequences with nonconservative nonsynonymous mutations (Figure **1c**) were advanced for further testing, and resynthesized to purge synonymous mutations. The purged sequences were then cloned into fresh p1 plasmid and platform cells, and cell growth was tested in the conditions used to evolve the mutant sequences and in which these unpurged sequences supported growth but the V124A mutant used as starting point did not. Unlike their unpurged counterparts, the purged sequences did not support growth, whereas the yeast THI4 positive control did so, as expected (Figure **1d**). The failure of the nonsynonymous mutations alone to confer improved growth indicates that the accompanying synonymous mutations were also necessary – or even sufficient – for the observed growth phenotype. An alternative explanation based on acquired beneficial nuclear mutations is *a priori* unlikely (i) because the genomic mutation rate is ∼100,000-fold lower than that of the OrthoRep target gene (Molina *et al*., 2022) and (ii) because we have seen no previous cases of this, let alone simultaneously in separate populations (García-García *et al*., 2022; Van Gelder et al., 2023).

That nonsynonymous mutations can improve the performance of a commercially codon-optimized bacterial enzyme in yeast is not surprising (Lanza *et al*., 2014). The codon-optimization algorithm strongly favored abundant yeast codons, which is not always the most effective scheme (Lanza *et al*., 2014), and indeed the synonymous mutations obtained all led to less-abundant codons, e.g., Gly GGT→GGC, Asp GAT→GAC. What *is* surprising is that the effects of synonymous mutations completely dominated the campaigns and that no new nonsynonymous mutations with substantial benefits were recovered. The nonsynonymous mutations were thus presumably near-neutral passenger mutations that were already present in genes in which beneficial synonymous mutations arose. Consistent with this possibility, the nonsynonymous mutations F14L and I198T (Figure **1c**) were previously classed as neutral or mildly deleterious (Van Gelder *et al*., 2023). Also, all but one of the eleven nonsynonymous mutations were at non-conserved or weakly conserved positions, and six of them occur naturally (Appendix **S2**). Note that OrthoRep’s extremely high mutation rate is designed to make many mutations in the target gene – this is a *feature*, not a bug – (Molina *et al*., 2022), so that patterns like those in Figure **1c** are sometimes to be expected.

A tactical conclusion from the failure to obtain mutant MhTHI4s substantially better than V124A or Y122C is that the mutational space accessible by TP-DNAP1_611 and TP-DNAP1_633, which make transition mutations but few transversions (García-García *et al*., 2022), has now been largely explored. Next-generation error-prone OrthoRep DNA polymerases that make many more transversions – and hence a far wider range of amino acid changes (Rix *et al*., 2023) – are likely to allow further progress.

A strategic conclusion for OrthoRep as an aerobic, eukaryotic platform to plantize non-plant enzymes is that users’ default assumption should be that synonymous mutations confer growth improvement as effectively as nonsynonymous ones – albeit just by increasing the enzyme’s expression level instead of changing its properties, i.e., by subverting the aim of the CDE campaign and creating an improvement mirage. To maximize efficiency, promising nonsynonymous mutations should therefore be tested without any accompanying synonymous mutations at an early stage, as in Figure **1d**. This strategic conclusion applies equally to using OrthoRep as a platform to improve plant enzymes because, whether the target enzyme gene is a native plant DNA sequence or a plant gene recoded for yeast expression, codon use could well be suboptimal, i.e., improvable by ‘yeastizing’ mutations.

To summarize: The codon bias issue illustrated here, like yeast’s preference for distinct amino acids at particular positions in a given protein (Van Gelder *et al*., 2024), must be monitored vigilantly when using OrthoRep as a platform to improve enzymes for use in plants. Codon and amino acid bias do not, however, compromise OrthoRep’s evolutionary power because yeastized features of the DNA or protein sequences they lead to can be swiftly recognized and rejected. Forewarned is forearmed.

## Author contributions

ADH, KVG, and AKG conceived and designed the project; KVG and AKG performed experiments; ADH carried out bioinformatic analyses and wrote the article, with contributions from other authors.

## Acknowledgments

We thank Dr. Chang C. Liu and his group members for their generous OrthoRep advice and help.

## References

García-García, J.D., Van Gelder, K., Joshi, J., Bathe, U., Leong, B.J., Bruner, S.D., Liu, C.C. et al. (2022) Using continuous directed evolution to improve enzymes for plant applications. Plant Physiol. 188, 971–983.

Gionfriddo, M., De Gara, L. and Loreto, F. (2019) Directed evolution of plant processes: Towards a green (r)evolution? Trends Plant Sci. 24, 999–1007.

Joshi, J., Li, Q., García-García, J.D., Leong, B.J., Hu, Y., Bruner, S.D. and Hanson, A.D. (2021) Structure and function of aerotolerant, multiple-turnover THI4 thiazole synthases. Biochem. J. 478, 3265–3279.

Lanza, A.M., Curran, K.A., Rey, L.G. and Alper, H.S. (2014) A condition-specific codon optimization approach for improved heterologous gene expression in Saccharomyces cerevisiae. BMC Syst. Biol. 8, 33.

Molina, R.S., Rix, G., Mengiste, A.A., Alvarez, B., Seo, D., Chen, H., Hurtado, J. et al. (2022) In vivo hypermutation and continuous evolution. Nat. Rev. Methods Primers 2, 37.

Ravikumar, A., Arzumanyan, G.A., Obadi, M.K.A., Javanpour, A.A. and Liu, C.C. (2018) Scalable, continuous evolution of genes at mutation rates above genomic error thresholds. Cell 175, 1946– 1957.e13.

Rix, G., Williams, R.L., Hu, V.J., Spinner, A., Pisera, A., Marks, D.S. and Liu, C.C. (2024) Continuous evolution of user-defined genes at 1 million times the genomic mutation rate. Science 386, eadm9073.

Van Gelder, K., Lindner, S.N, Hanson, A.D. and Zhou, J. (2024) Strangers in a foreign land: ‘Yeastizing’ plant enzymes. Microb. Biotechnol. 17, 14525.

Van Gelder, K., Oliveira-Filho, E.R., García-García, J.D., Hu, Y., Bruner, S.D. and Hanson, A.D. (2023) Directed evolution of aerotolerance in sulfide-dependent thiazole synthases. ACS Synth. Biol. 12, 963–970.

Zhong, Z., Ravikumar, A. and Liu, C.C. (2018) Tunable expression systems for orthogonal DNA replication. ACS Synth. Biol. 7, 2930–2934.

