## Supporting information for "Mirages in continuous directed enzyme evolution: A cautionary case study with plantized bacterial THI4 enzymes"

### Experimental procedures

*Chemicals and media*

Drop-out mix minus histidine, leucine, tryptophan, uracil without yeast nitrogen base and yeast nitrogen base without amino acids, ammonium sulfate, and thiamin were from US Biologicals (Salem, MA). Zymolyase was from AMSBio (Cambridge, MA). Phusion PCR, proteinase K, and RNase A were from Thermo Scientific (Waltham, MA). G418 sulfate was from Santa Cruz Biotechnology Inc. (Santa Cruz, CA). Restriction enzymes and site-directed mutagenesis kits were from New England Biolabs (Ipswich, MA). All other chemicals were from US Biologicals or Fisher Scientific.

*THI4 sequences and OrthoRep constructs*

MhTHI4 sequences were derived from the codon-optimized gene used previously (Van Gelder *et al*., 2023). For evolution campaigns, the mutant Mh_V124A sequence (Van Gelder *et al*., 2023) was amplified from OrthoRep cell DNA, digested with NsiI and SphI, and ligated into GR-306MP containing a 72A (full-length), 24A, or 0A poly(A) tail. The tail was shortened or removed by site-directed mutagenesis using the 72A construct as template and primers in Table **S2**. Constructs were transformed into strain GA-Y319 containing p1 and p2 as before (Van Gelder *et al*., 2023). GA-Y319 harboring p1 containing Mh_V124A was the donor strain in protoplast fusions with recipient strain BY4741 *his3*∆ *leu2*∆ *met15*∆ *ura3*∆ *thi4*∆ containing nuclear plasmid ArEC bearing TP-DNAP1_611 or TP-DNAP1_633 (García-García *et al*., 2022). GA-Y319 cells harboring p1_ Mh_V124A plasmids were grown in synthetic complete medium (SC) -Leu. BY4741 *thi4*∆ cells harboring ArEC were grown in SC -His -Trp and BY4741 *thi4*∆ harboring ArEC and p1_Mh_V124A plasmids were grown in SC -His -Trp -Leu.

*Evolution campaigns*

Campaigns were run using 3-mL cultures in 24-well plates from MilliporeSigma (St. Louis, MO) in a Multitron HT 3 mm platform shaker (INFORS HT, Annapolis Junction, MD) at 30°C, 800 rpm, and 80% relative humidity. Cultures were started at an $OD_{600}$ of 0.05 and washed thrice with thiamin- and 5-(2-hydroxyethyl)-4-methylthiazole (HET)-free medium before transfer from thiamin medium to thiamin-free or HET-free medium. Cultures grew for 4-6 days in each passage; passage length decreased as campaigns proceeded and cultures reached a higher OD faster. In the cold turkey selection strategy, cultures were grown from the start without added thiamin or HET. In the gradual selection strategy, cultures were first given limiting (10 nM) thiamin, which allowed them to reach an $OD_{600}$ of 1, then switched to HET supplementation, beginning with a luxury concentration (300 nM), then moving to successively lower concentrations (30, 10, and 3 nM) and finally to HET-free medium. The rationale for tapering with HET rather than thiamin is that thiamin represses the whole thiamin synthesis pathway in yeast (Kowalska and Kozik, 2008) but HET does not (Praekelt *et al*., 1994). HET is thus predicted to favor survival of the transition from low to zero supplementation as previously HET-supplemented cells can make thiamin at once after transfer whereas previously thiamin-supplemented cells cannot and so enter a thiamin-less 'valley of death'. Tests with cells transferred from low-thiamin to zero-thiamin medium confirmed this prediction.

*Bacterial contamination checks*

During evolution campaigns, cultures were periodically checked for bacterial contamination by (i) spot-plating cultures on LB medium and incubating at 30°C for 1-3 days, or (ii) PCR-amplifying the 16S bacterial ribosomal DNA from DNA isolated from yeast cultures using the primers given in Table **S2**. Note that the G418 antibiotic used in OrthoRep protocols (García-García *et al*., 2022) did not prevent or eliminate bacterial contamination. Contaminated cases were replaced from uncontaminated stocks.

*DNA extraction, sequencing, and synthesis*

Genomic DNA was extracted from cultures as described (García-García *et al*., 2022) for analysis of p1 plasmids by gel electrophoresis, or as follows for sequencing or contamination checks: 100-300 µL of culture was centrifuged (15,000 × *g*, 3 min) and resuspended in 200 mM lithium acetate and 1% SDS and incubated for 5 min at 70°C. Three hundred µL of 96-100% ethanol was added and vortex-mixed for 5 s. After centrifuging as above, the pellet was washed with 100 µL of 70% ethanol, resuspended in 100 µL of water or TE (10 mM Tris-Cl, pH 7.4, 1 mM $Na_2$EDTA), and centrifuged (15,000 × *g*, 15 s). The supernatant was transferred to a fresh tube; 1 µL was used in PCR reactions. Sequencing was done by Azenta (Burlington, MA) using primers in Table **S2**. Sequences were analyzed with Benchling (San Francisco, CA). Sequences without synonymous mutations were generated by site-directed mutagenesis using primers in Table **S2** or synthesized by Twist Biosciences (San Francisco, CA).

*Testing of evolved sequences*

Mutant sequences (purged of synonymous mutations) were PCR-amplified with primers from Table **S2**, digested with NsiI and SphI, ligated into GR-306MP with a 72A tail, and cloned into BY4741 *his3*Δ *leu2*Δ *met15*Δ *ura3*Δ *thi4*Δ containing ArEC_611 as above. Wildtype yeast THI4 (ScTHI4) and an empty vector control were cloned into BY4741 *his3*Δ *leu2*Δ *met15*Δ *ura3*Δ *thi4*Δ containing ArEC_611 as benchmarks for the growth assays with the new mutant sequences and the parent V124A sequence. Twelve clones of each mutant and each control were grown in 3-mL cultures of SC -Leu -His -Trp with 300 nM thiamin for two days. Cultures were washed thrice with SC medium without thiamin and inoculated into fresh 3-mL cultures of SC -Leu -His -Trp media without thiamin at $OD_{600}$ of 0.05. Cultures were grown for 13 days and $OD_{600}$ was monitored.

### References for experimental procedures

García-García, J.D., Van Gelder, K., Joshi, J., Bathe, U., Leong, B.J., Bruner, S.D., Liu, C.C. *et al.* (2022) Using continuous directed evolution to improve enzymes for plant applications. *Plant Physiol.* **188**, 971–983.

Kowalska, E. and Kozik, A. (2008) The genes and enzymes involved in the biosynthesis of thiamin and thiamin diphosphate in yeasts. *Cell. Mol. Biol. Lett.* **13**, 271–282.

Praekelt, U.M., Byrne, K.L. and Meacock, P.A. (1994) Regulation of THI4 (MOL1), a thiamine-biosynthetic gene of *Saccharomyces cerevisiae*. *Yeast* **10**, 481–490.

Van Gelder, K., Oliveira-Filho, E.R., García-García, J.D., Hu, Y., Bruner, S.D. and Hanson, A.D. (2023) Directed evolution of aerotolerance in sulfide-dependent thiazole synthases. *ACS Synth. Biol.* **12**, 963–970.

**Table S1.** Conservation of MhTHI4 V124A mutated residues among representative prokaryotic THI4s

Diverse THI4 sequences from 199 genomes (Joshi *et al.*, 2021) were aligned and compared to the nonsynonymous mutations in MhTHI4. Natural replacements corresponding to mutations are in red.

| **Mutation** | **Natural replacements** |
|---|---|
| F14L | **L** Y Q A H M S I T K V N |
| I27T | V L F |
| L35S | I M F V |
| Y40H | **H** K A E F D R V N T |
| T117A | **A** R K V M H L Q S N E I |
| Y122C | T S F M V I L W A |
| A124V | **V** I M F L |
| V127A | **A** I L |
| I198T | V M A C P S I |
| E202G | D |
| K244E | **E** Q R L A |

**Table S2.** Primers for cloning, sequencing, and site-directed mutagenesis

Restriction sites are underlined. Primers used for cloning into GR-306MP contain an additional CAT sequence (**bold**, encoding histidine) after the start codon, to create a NsiI restriction site.

| Primer name | Sequence (5′–3′) | Purpose |
|---|---|---|
| MhTHI4_NsiI_F | CGATATG**CAT**GAAAAGATTGTTTCTGCTGGTATTG | Cloning of *MhTHI4* into GR-306MP |
| MhTHI4_SphI_R | ATCGGCATGCTTACAATGAATCACAAATCATTTGTGC | Cloning of *MhTHI4* into GR-306MP |
| ScTHI4_NsiI_F | CGATATG**CAT**TCTGCTACCTCTACTGCTA | Cloning of *ScTHI4* into GR-306MP |
| ScTHI4_SphI_R | ATCGGCATGCCTAAGCAGCAAAGTGTTTC | Cloning of *ScTHI4* into GR-306MP |
| p1_F | TTATTGGAAGATTAGTACGTCTCC | Sequencing *MhTHI4, ScTHI4* in p1 |
| Leu_Short_R | GCTGTGATTTCTTGACCAACGTGG | Sequencing *MhTHI4, ScTHI4* in p1 |
| MhTHI4_Int_F | GCACCAGTTATTAGAGAGGGTATG | Sequencing *MhTHI4* in GR-306MP and p1, SDM of GR-306MP_*MhTHI4*_Y122C |
| MhTHI4_Int_R | GCTTTAGCCAAGTAATATGCAGC | Sequencing *MhTHI4* in GR-306MP and p1 |
| ScTHI4_Int_F | CGGTATGAAGGGTCTGGACATGAACC | Sequencing of *ScTHI4* in GR-306mP and p1 |
| ScTHI4_Int_R | GGTTCATGTCCAGACCCTTCATACCG | Sequencing of *ScTHI4* in GR-306mP and p1 |
| GR-306MP_0A_F | GGGAATTGGGATGTCATGCTTTT | Constructing 0A GR-306MP from 72A GR-306MP by SDM |
| GR-306MP_0A_R | TCATGGGGCATCGCATGCTTATCTGTG | Constructing 0A GR-306MP from 72A GR-306MP by SDM |
| GR-306MP_24A_F | CCTGTCACCGGATGTGTT | Constructing 24A GR-306MP from 72A GR-306MP by SDM |
| GR-306MP_24A_R | TTTTTTTTTTTTTTTTTTTTTTTTCATGGGGCATC | Constructing 24A GR-306MP from 72A GR-306MP by SDM |
| K244E_F | GTTATCTGGTGAAAAGGCTGCAC | SDM of GR-306MP_*MhTHI4*_V124A |
| K244E_R | AACATACCACCAAAAATTGGAC | SDM of GR-306MP_*MhTHI4*_V124A |
| L137S_R | CCAATTAACAACTGAACCAGCAACT | SDM of GR-306MP_*MhTHI4*_Y122C |
| V127A_F | TGTTGCTCAGTTGAAGATGCTGTT | SDM of GR-306MP_*MhTHI4*_Y122C |

| | | |
|---|---|---|
| V127A_R | ATTGAAAATTGTTGCACCAGCTCTAG | SDM of GR-306MP_*MhTHI4*_Y122C |
| 27_F | AGAGTTTGATCMTGGCTCAG | Amplification of 16S ribosome in Betaproteobacteria |
| 1492_R | TACGGYTACCTTGTTACGACTT | Amplification of 16S ribosome in Betaproteobacteria |
| 63_F | CAGGCCTAACACATGCAAGTC | Amplification of 16S ribosome in Alphaproteobacteria |
| M1387_R | GGGCGGWGTGTACAAGRC | Amplification of 16S ribosome in Alphaproteobacteria |
